# An experimental study of free flight kinematics in a miniature parasitoid wasp *Trichogramma telengai*

**DOI:** 10.1101/2024.03.21.586056

**Authors:** Nadezhda A. Lapina, Sergey E. Farisenkov, Evgeny O. Shcherbakov, Dmitry Kolomenskiy, Alexey A. Polilov

## Abstract

Body size is the major factor to the flight mechanics in animals. To fly at low Reynolds numbers, miniature insects have adaptations in kinematics and wing structure. Many microinsects have bristled wings, which reduce inertia and power requirements when providing good aerodynamic efficiency. But both bristled and membranous-winged microinsects fly at Reynolds numbers of about 10. Yet, the kinematics of the smallest known membranous-winged species have not been studied sufficiently. The available data are limited to the forewings of a relatively large parasitoid wasp *Encarsia formosa*. We studied kinematics of wings and body and flight performance in one of the smallest membranous-winged wasps, Trichogramma telengai (0.5 mm body length, *Re* = 12). *T. telengai* reaches 29 cm s^-1^ speed and 7 m s^-2^ acceleration in horizontal flight which are comparable with the flight performance of other microinsects. The wingbeat cycle is characterized by high frequency (283 Hz) and stroke amplitude (149°) and includes U-shaped strokes at high angles of attack and prolonged clap-and-fling. The hindwings move with a slight phase shift and smaller amplitude than the forewings. *T. telengai* differs from large membranous-winged insects and miniature featherwing beetles in kinematics, but it is fundamentally similar to *E. formosa* (*Re* = 18, membranous wings) and thrips *Frankliniella occidentalis* (*Re* = 15, bristled wings). We showed that, at *Re* ≈ 10^1^, both membranous and bristled-winged insects have sufficient flight performance. Further study of the bristled-winged insects will make it possible to define the size limits of effectiveness of different wing structures.

## Introduction

Insects vary in size in a much wider range than any class of vertebrates (Polilov, 2016a; Vendl and Šípek, 2016). The adults of the smallest insects are 3 orders of magnitude smaller in body length, and the mass is 7 orders of magnitude less, compared to the largest insects. The large and the small insects alike share the ability to fly, but the underlying aerodynamic mechanisms change with the size. To stay aloft, a flying insect must generate an upward force equal in magnitude to its body weight. The flapping wings press against the air and deflect the air flow momentum downwards. In many large insects with membranous wings, the trajectory of the wingtip is almost planar. The aerodynamic force is perpendicular to the stroke plane as it is dominated by the Joukowski lift, which is an inviscid mechanism, i.e., the force is due to the inertia of air.

As long as we consider smaller insects, the mass, to which the inertial force is proportional, decreases in proportion to the cube of the representative linear dimension. The surface area, to which the friction force is proportional, decreases as the square of the linear dimension. This means that the ratio of friction force to inertia force increases (Brodsky, 1988; Sane, 2003; Walker, 2003). The role of the viscous friction becomes essential for the insects that are less than 1 mm long. Under such conditions, wings moving in a horizontal plane at low angles of attack can no longer generate sufficient vertical force. Therefore, the insect is forced to increase the angle of attack of the wing and change the trajectory of its movement so that, during the power stroke, the drag force of the wing is oriented approximately upward. As miniaturization progresses, the trajectory of the wing apex during translational phases deforms in the shape of the letter U, and the average angle of attack increases (Sun, 2023; Kolomenskiy, 2023).

A flapping wing cyclically changes the direction of its path. Conceptually, more energy is required to accelerate the wing after each reversal, as compared with unilateral translation. But insects evolved to use unsteady aerodynamics to mitigate the negative effects of reciprocating motion. One of those unsteady mechanisms is known as the clap-and-fling. At stroke reversals, the two wings press together and open again such that the costal veins separate first and the membrane follows. The clap-and-fling allows insects to maximize the wing beat amplitude and make use of the aerodynamic interference. The miniature insects adapted the clap-and-fling trajectory with a large vertical excursion: the wings are pressed together until they reach the most elevated point of the U-shaped trajectory, then the wings fling apart, sideways and downwards (Cheng and Sun, 2019).

Many of the miniature insects such as parasitoid wasps, thrips and Ptiliidae beetles have bristled wings (Polilov, 2016), which are an important adaptation to reduce mechanical power required for flight at low Reynolds numbers (Farisenkov et al., 2022). However, there are a lot of miniature membranous-winged insects, including one of the most well-studied species in terms of wing kinematics, *Encarsia formosa* (Hymenoptera; Aphelinidae), with the body length of approximately 0.69 mm (Singh and Sood, 2018). Torkel Weis-Fogh first described the clap-and-fling in that species (Weis-Fogh, 1973). Since then, the flight mechanics of *E. formosa* have been studied in detail (Cheng and Sun, 2016; Cheng and Sun, 2018).

In this work, we consider the kinematics of one of the smallest membranous-winged wasps *Trichogramma telengai* (Hymenoptera: Trichogrammatidae) with a body length of 0.48 mm. The movements of not only the forewings, but also the hindwings and body are reconstructed. Obtained data allow us to compare the adaptations to flight at the miniature scale between different taxonomic groups of Hymenoptera.

## Materials and methods

### Insects

This study was carried out on a parthenogenetic laboratory line of *Trichogramma telengai* Sorokina, 1987 (Hymenoptera: Trichogrammatidae). The line was cultivated in Sergey Reznik’s Laboratory of Experimental Entomology, Zoological Institute, Russian Academy of Sciences, St. Petersburg, Russia, during many years under constant laboratory conditions on eggs of the grain moth *Sitotroga cerealella* (Lepidoptera: Gelechiidae) (Reznik and Voinovich, 2019).

### Morphology and morphometrics

To measure live body mass we immobilized 404 specimens with carbon dioxide and weighed them on Sartogosm МВ210-А (Sartogosm LLC, Russia) analytical balance. Part of the sample was fixed in 70% ethanol for morphological study. Body length was measured on high-speed video recordings as triangulated distance between the end of the abdomen and the point in-between the antennal bases.

For wing morphometrics, 15 forewinds and 15 hindwinds were dissected from the body. Before dissection insects were softened for 24 hours in a 5:5:1 mixture of ethanol, glycerol and water. Then wings were washed in 100% ethanol twice for 24 hours and placed into Eukitt UV (Orsatec) medium for 24 hours. After that we made permanent slide preparations in Eukitt UV resin. The preparations were imaged on Olympus BX43 (Olympus) light microscope. We measured wing length, area, radius of gyration and percentage of wing surface area occupied by setae from digital photographs (Fig. 1).

**Figure 1.**
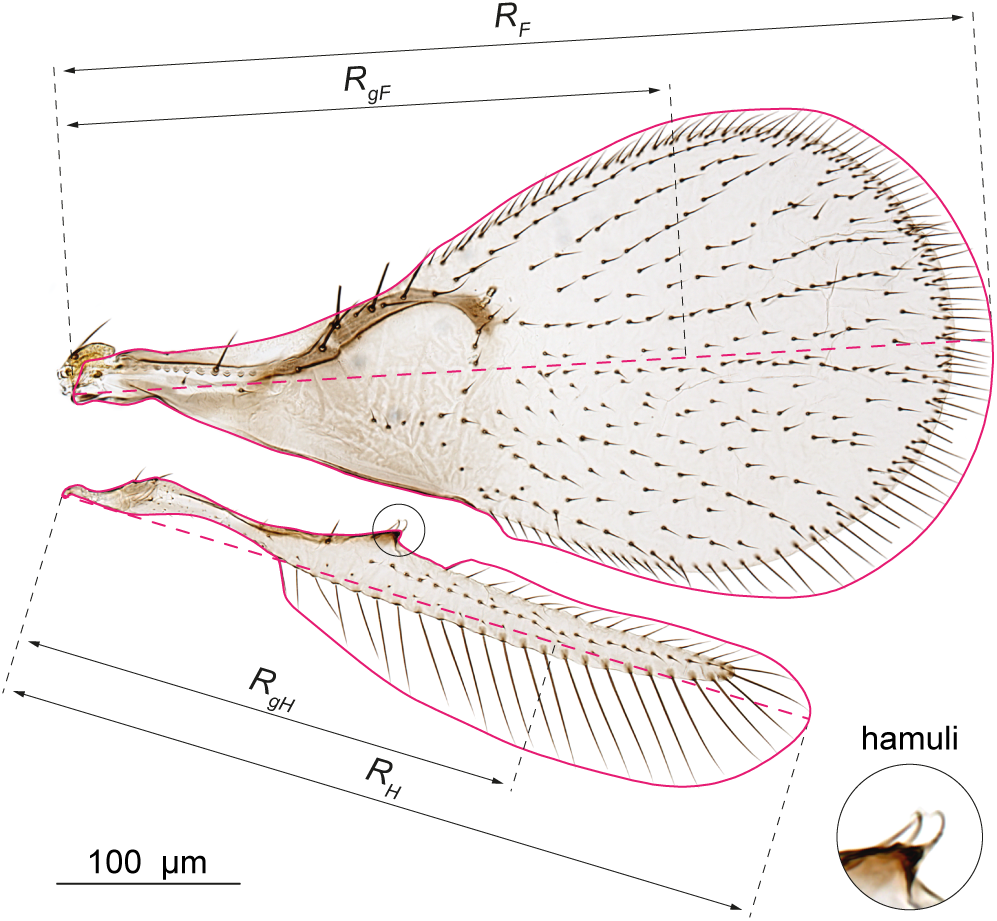
*Trichogramma telengai* wings morphometrics scheme. *R_F_* – forewing length; *R_H_* –hindwing length; *R_gF_*– forewing radius of gyration; *R_gH_* – hindwing radius of gyration; red outline – region of wings area measurement.

### High-speed recording

Free flight of *T. telengai* was recorded with a frequency of 4600 FPS in transmitted infrared 850 nm LED light. Flight chamber was additionally illuminated by white LED to attain the illumination level natural for the insects. Four Evercam 4000 (Evercam, Russia) high speed cameras are placed radially symmetrical around the horizontal axis at an angle of 60° to each other (Fig 2A), their optical axes intersect at the focal point of the lenses. We used 25 mm f/2.8 2.5-5X Ultra Macro lenses (Laowa) with a low level of distortion. About 30 specimens were simultaneously placed in a closed flight chamber of complex shape (volume 3.8 cm^3^, distance between the glasses, through which the recording was performed, 15 mm). During the recording, the flight chamber temperature was stabilized in the range of 25 to 27°С by air cooling.

**Figure 2.**
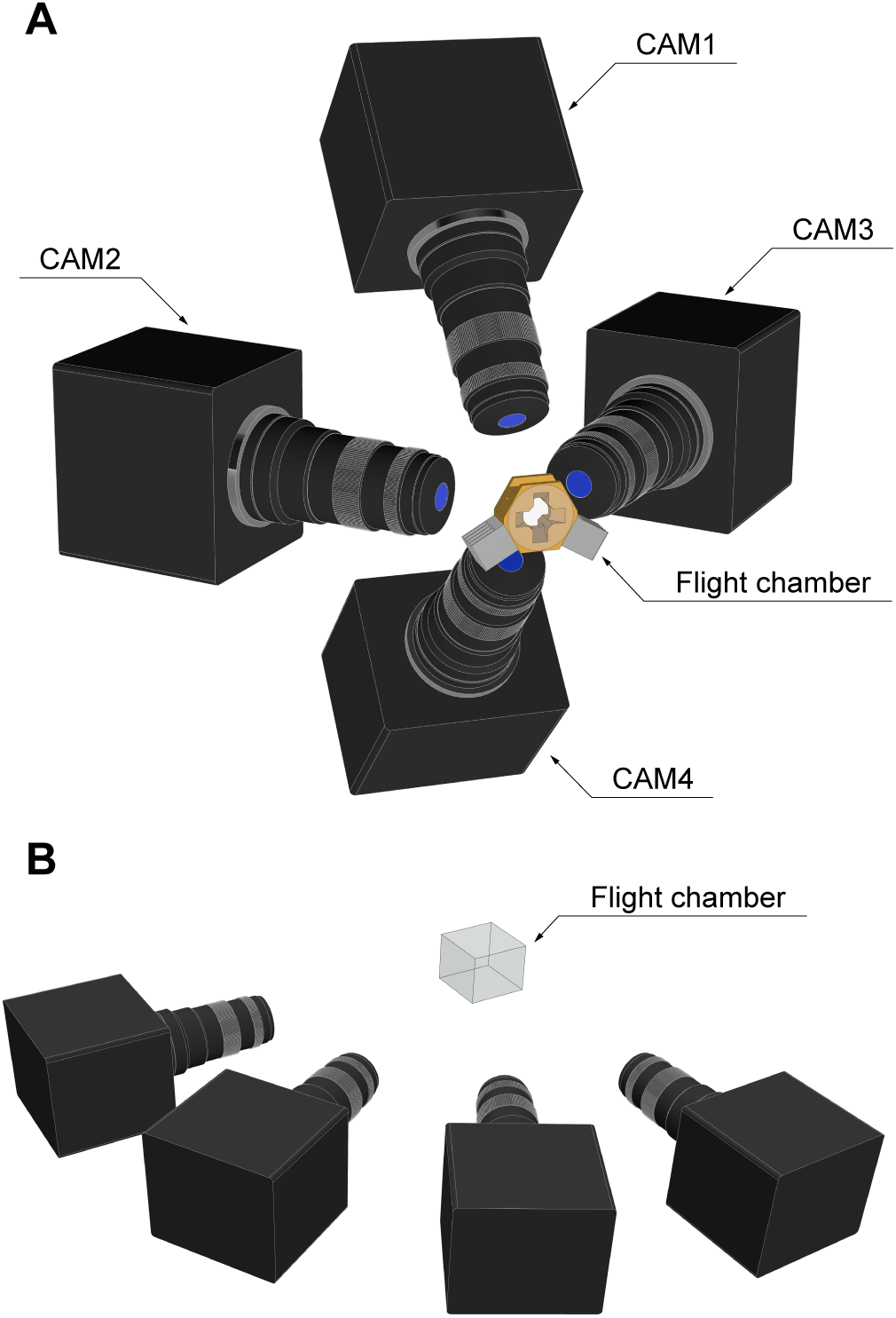
Schemes of high-speed video recording setups. (A) Macro filming for kinematics reconstruction. (B) Recording of flight trajectories in a spacious chamber.

For the analysis of flight performance we filmed insects with a frequency of 200 FPS in a spacious glass chamber of 58×58×47 mm size in transmitted infrared and ambient white light. The width of the chamber is at least 120 times larger than the length of *T. telengai*, allowing insects to fly freely. In this setup, the cameras were placed horizontally (Fig. 2B).

### Tracking and triangulation

For analysis of the kinematics, we selected 6 recordings with the most linear flight in focus on all 4 cameras. In cases where more than 4 strokes were recorded under these conditions, the recording was subdivided to avoid averaging different flight modes. We analyzed 11 recording fragments of 3-4 consecutive strokes, 36 strokes in total.

2D trajectories of 18 landmarks on wings and body were obtained in Tracker (Open Source Physics) on each camera projection (Fig. 3A). For triangulation we used DLTdv8a software (Hedrick, 2008). The cameras were calibrated by a 2×2 mm dot grid with a step of 0.25 mm.

**Figure 3.**
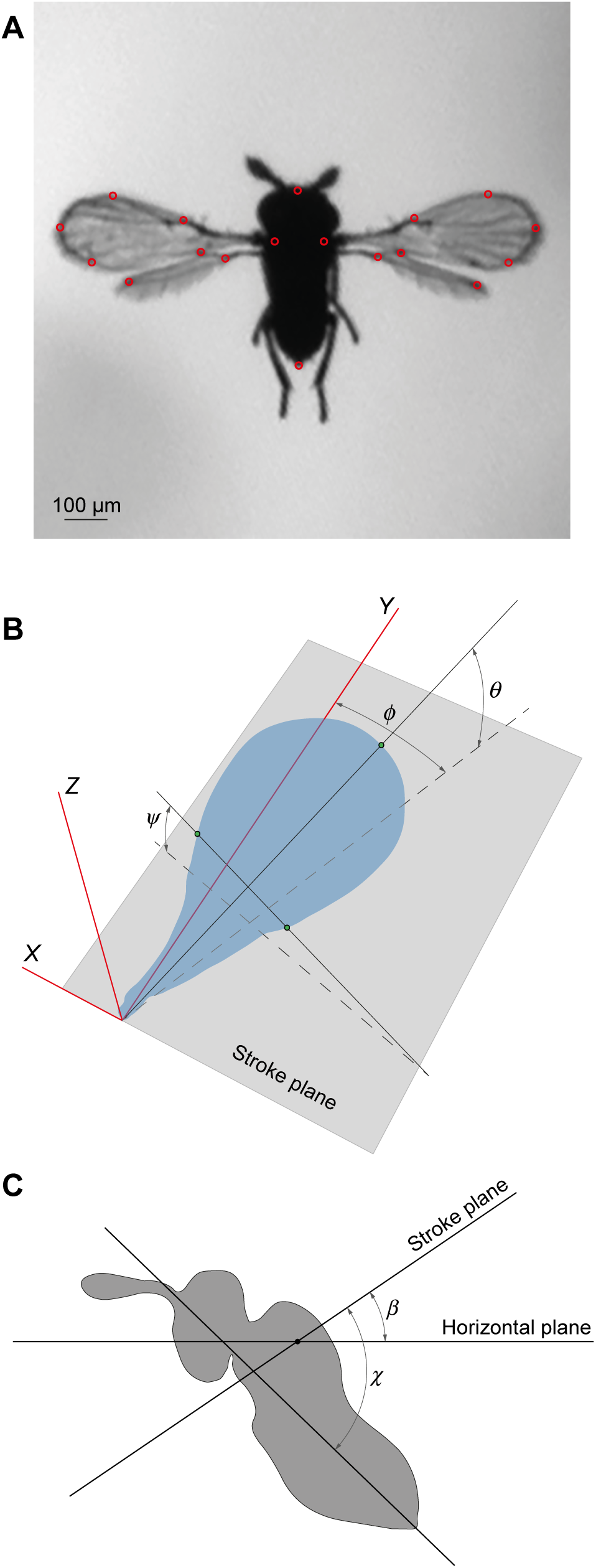
3D reconstruction of *T. telengai* flight kinematics. (A) Location of landmarks on wings and body. (B) Measurement scheme for Euler angles. (C) Measurement scheme for angle of body pitch (*χ*) and stroke plane inclination angle relative to the laboratory horizontal plane. (*β*).

Similarly, we obtained the trajectories of the insects in the large flight chamber on 4 camera projections with Tracker and triangulated them in DLTdv. 25 trajectories with overall flight time of 16.0 s were reconstructed on 9 records.

### Three-dimensional reconstruction and analysis of flight kinematics and trajectory

For the mathematical description of the wings kinematics, we used the Euler angles system (Fig. 3B): stroke amplitude (*φ*), stroke deviation angle (*θ*) and pitch angle (*ψ*) (Cheng and Sun, 2016). We also calculated the angle of attack (*α*) - the angle between the wing plane and its velocity vector. The Euler angles and angle of attack were calculated in the coordinate system associated with the body and the stroke plane: the origin is between the bases of the forewings, the *XOY* plane is a plane parallel to the stroke plane, and the *XOZ* plane is a plane parallel to the sagittal plane. The stroke plane and the wing plane were approximated using all wing landmarks for the current stroke. We defined the start of every cycle (*t/T* = 0) as the moment when the forewing angle *φ* reaches its minimal value during the supination. After calculating the Euler angles and angle of attack for each frame in the recording fragment, we calculate the mean values for left and right wings, separated the strokes and calculate their means over all cycles. For the forewings and hindwings we calculated all angles separately. *Re* was calculated as follows:

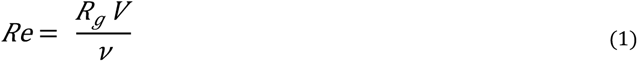

Where *R_g_*is the radius of gyration of the forewing, *V* is wing velocity at the radius of gyration and *ν* is the kinematic viscosity of air at 25°С.

To perform comparative analysis we recalculated *Re* for *Encarsia formosa* and *Frankliniella occidentalis* using this formula, because available *Re* values are based on wing chord as characteristic length (Lyu et.al., 2019).

Body orientation was described by two angles in the laboratory coordinate system: the stroke plane pitch angle (*β*) relative to horizon; the body pitch angle (*χ*) as the angle of the body axis relative to the stroke plane (Fig. 3C).

Horizontal and vertical components of speed and acceleration were calculated in the laboratory coordinate system based on the trajectory of the point in-between the forewing bases landmarks. Average stroke frequency was calculated as the median of all recordings by counting the number of frames in several cycles for each recording (24 recordings, 242 cycles in total). All calculations were automated with a Matlab (Mathworks) script.

Mean and maximum speeds and accelerations within the larger flight chamber were calculated using the values from all tracks. Three-dimensional coordinates obtained in DLTdv were smoothed with local polynomial regression, using function loess of the package stats in R (R Core Team). Smoothed coordinates were used to calculate flight velocity between the neighboring trajectory points. These values of velocity were filtered using a step = 3 moving average and the accelerations were calculated based on these final values. The average values were calculated as the medians owing to the non-normal distribution of the respective samples, while the 99th percentiles were taken as the maximum values to exclude the influence of outliers. We also separately analyzed velocities and accelerations in horizontal flight, selecting the segments of trajectories with the deviation from the plane of horizon of no more than 30°.

## Results

The mean body mass of *Trichogramma telengai* is 6.1 µg, the body lengths of recorded specimens range from 0.40 to 0.50 mm with the median of 0.48 mm. Bristles occupy 24.4% of the total wing area (Table 1).

**Table 1.**
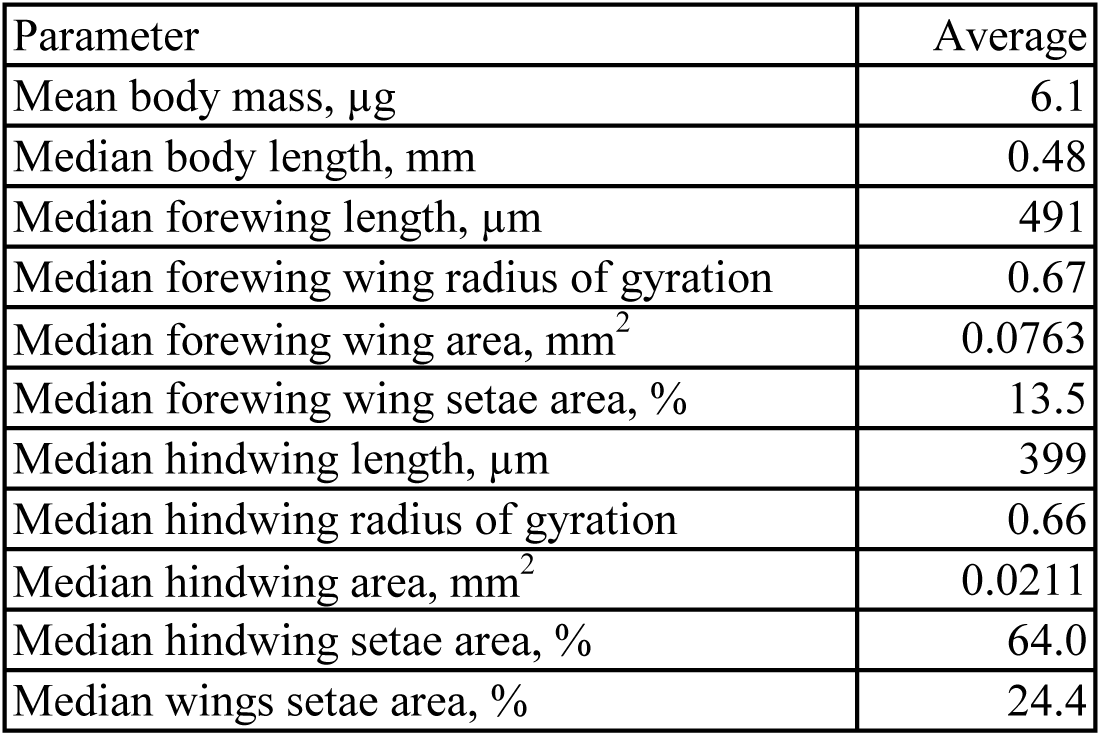
Morphological parameters of *Trichogramma telengai*.

| Parameter | Average |
| --- | --- |
| Mean body mass, µg | 6.1 |
| Median body length, mm | 0.48 |
| Median forewing length, µm | 491 |
| Median forewing wing radius of gyration | 0.67 |
| Median forewing wing area, mm<sup>2</sup> | 0.0763 |
| Median forewing wing setae area, % | 13.5 |
| Median hindwing length, µm | 399 |
| Median hindwing radius of gyration | 0.66 |
| Median hindwing area, mm<sup>2</sup> | 0.0211 |
| Median hindwing setae area, % | 64.0 |
| Median wings setae area, % | 24.4 |

We calculated main kinematic parameters in 11 video sequences (TT1 - TT10) (SI Video 1). The values for each are presented in table 2. The flight speed of recorded specimens in the small chamber is quite slow (from 5.2 to 17.7 cm s^-1^) and advance ratio *J* (body velocity divided by mean wingtip speed) is in the range between 0.06 and 0.20. Thus, 4 of 11 video sequences represent hovering, according to the advance ratio threshold 0.1 proposed by Ellington as the definition of hovering flight (Ellington, 1984). *T. telengai* fly significantly faster in the spacious flight chamber than in the small one: the mean and the maximal flight speed are, respectively 12.2 and 32.7 cm s^-1^, when speeds in horizontal flight are 13.1 and 29.1 cm s^-1^ respectively. Maximal horizontal acceleration is 7.1 m s^-2^. Mean speed in horizontal flight is slightly greater than mean speed in general because a significant part of the tracks represents flight from bottom to the top of the flight chamber.

**Table 2.** Kinematic parameters of *Trichogramma telengai* specimens TT1-TT11.

| Parameter | Median | TT1 | TT2 | TT3 | TT4 | TT5 | TT6 | TT7 | TT8 | TT9 | TT10 | TT11 |
| --- | --- | --- | --- | --- | --- | --- | --- | --- | --- | --- | --- | --- |
| Wingbeat frequency, Hz | 290 | 293 | 297 | 261 | 290 | 288 | 290 | 290 | 290 | 293 | 287 | 292 |
| $Re$ | 12.5 | 10.0 | 10.1 | 12.1 | 12.5 | 12.5 | 12.7 | 12.7 | 13.0 | 13.3 | 10.2 | 12.1 |
| Flight speed horizontal, $m\ s^{-1}$ | | 0.078 | 0.101 | 0.152 | 0.173 | 0.090 | 0.063 | 0.022 | 0.067 | 0.093 | 0.134 | 0.154 |
| Flight speed vertical, $m\ s^{-1}$ | | 0.032 | 0.029 | -0.035 | 0.000 | 0.027 | 0.037 | 0.047 | 0.043 | 0.049 | -0.040 | -0.087 |
| Flight speed total, $m\ s^{-1}$ | | 0.084 | 0.106 | 0.156 | 0.173 | 0.094 | 0.073 | 0.052 | 0.080 | 0.106 | 0.140 | 0.177 |
| Advance ratio ($J$) | | 0.10 | 0.13 | 0.18 | 0.19 | 0.11 | 0.08 | 0.06 | 0.09 | 0.11 | 0.18 | 0.20 |
| Acceleration horizontal, $m\ s^{-2}$ | | 2.75 | 2.58 | 5.49 | -2.73 | -6.22 | -2.90 | 5.11 | 5.22 | 0.44 | -5.29 | -7.30 |
| Acceleration vertical, $m\ s^{-2}$ | | -0.29 | 0.05 | -0.43 | 2.61 | -0.10 | 3.58 | -0.57 | -5.08 | 3.18 | 2.68 | 0.36 |
| Stroke plane pitch angle ($\beta$), deg | | 3.5 | 2.0 | 32.8 | -9.7 | -14.3 | -25.0 | -20.9 | -12.5 | -13.2 | -27.3 | -5.7 |
| Body pitch angle ($\chi$), deg | | 52.6 | 51.1 | 50.2 | 52.5 | 51.5 | 48.3 | 47.4 | 48.9 | 49.8 | 51.0 | 53.6 |
| Body pitch amplitude ($\chi$), deg | 12.3 | 14.1 | 10.0 | 12.3 | 9.9 | 9.7 | 13.8 | 12.5 | 12.4 | 12.5 | 6.6 | 9.0 |
| Body to horizon pitch angle (χ-β), deg | | 49.0 | 49.1 | 17.3 | 62.2 | 65.8 | 73.3 | 68.3 | 61.4 | 63.0 | 78.3 | 59.3 |
| Upstroke duration, $t/T$ | 0.25 | 0.257 | 0.256 | 0.244 | 0.27 | 0.265 | 0.243 | 0.253 | 0.247 | 0.246 | 0.254 | 0.272 |
| Up-moving phase duration, $t/T$ | 0.26 | 0.258 | 0.245 | 0.299 | 0.23 | 0.257 | 0.266 | 0.247 | 0.265 | 0.248 | 0.27 | 0.257 |
| Downstroke duration, $t/T$ | 0.37 | 0.367 | 0.384 | 0.356 | 0.397 | 0.387 | 0.349 | 0.368 | 0.352 | 0.367 | 0.379 | 0.376 |
| Forewing stroke amplitude ($\Delta\phi$), deg | 149.0 | 144.3 | 145.9 | 147.8 | 147.2 | 149.4 | 149.3 | 149.0 | 152.0 | 152.2 | 147.1 | 150.6 |
| Forewing downstroke angle of attack max, deg | 71.8 | 61.7 | 60.4 | 55.2 | 64.6 | 66.8 | 82.9 | 80.9 | 74.5 | 75.9 | 77.3 | 71.8 |
| Forewing upstroke angle of attack max, deg | 75.3 | 78.9 | 74.5 | 75.3 | 90.1 | 82.5 | 89.0 | 72.9 | 71.6 | 72.1 | 73.6 | 89.7 |
| Forewing downstroke deviation angle ($\theta$) min, deg | 4 | 3.4 | 4.1 | 2.8 | 6.1 | 5.1 | 5.4 | 3.6 | 3.6 | 4.2 | 3.5 | 3.4 |
| Forewing downstroke deviation angle ($\theta$) max, deg | 52 | 50.5 | 50.8 | 53.8 | 48.9 | 51.1 | 60.5 | 59.0 | 58.9 | 60.3 | 50.4 | 52.3 |
| Forewing upstroke deviation angle ($\theta$) min, deg | -20 | -20.6 | -20.4 | -19.8 | -16.6 | -18.8 | -19.1 | -20.2 | -23.0 | -23.3 | -16.4 | -16.9 |
| Forewing upstroke deviation angle ($\theta$) max, deg | 33 | 32.6 | 32.2 | 30.4 | 36.3 | 34.2 | 34.3 | 28.8 | 30.1 | 28.7 | 35.8 | 36.6 |
| Hindwing stroke amplitude ($\Delta\phi$), deg | 102 | 99.4 | 99.2 | 98.8 | 102.1 | 105.1 | 110.8 | 106.2 | 109.3 | 107.5 | 102.0 | 98.9 |
| Hindwing downstroke deviation angle ($\theta$) min, deg | 0 | -2.2 | -0.7 | -0.3 | 2.3 | 1.2 | 1.7 | 1.0 | -1.3 | 1.0 | -1.2 | -0.7 |
| Hindwing downstroke deviation angle ($\theta$) max, deg | 34 | 31.1 | 33.0 | 39.5 | 34.5 | 33.7 | 42.5 | 40.6 | 41.2 | 42.3 | 31.7 | 33.6 |
| Hindwing upstroke deviation angle ($\theta$) min, deg | -26 | -26.3 | -27.0 | -26.2 | -22.8 | -23.8 | -23.0 | -27.2 | -28.9 | -25.6 | -24.4 | -21.9 |
| Hindwing upstroke deviation angle ($\theta$) max, deg | 14 | 11.8 | 16.4 | 13.6 | 19.5 | 11.7 | 19.1 | 14.4 | 13.3 | 11.1 | 15.7 | 16.9 |

During the U-shaped downstroke (*t/T* ≈ 0.58 – 0.95, hereinafter median values) and upstroke (*t/T* ≈ 0.06 – 0.33) the forewings move at high angles of attack up to 72° and 75° respectively (Fig. 4A). Wings stroke at amplitude 149° and clap above the body. The clap-and-fling motion is prolonged because of the presence of the recovery stroke (*t/T* ≈ 0.33 –0.58). This phase was noted first time in *E. formosa* as “moving up” and was defined as a relatively slow upward movement of wings after closing and before fling (Cheng and Sun, 2021). With a mean wingbeat frequency of 283 Hz, the cycle-averaged mean Reynolds number is 12. *Re* reaches 24 during upstroke when the wing speed is maximal (1.5 m s^-1^) (Fig. 4B). Downstroke is much slower (1.1 m s^-1^).

**Figure 4.**
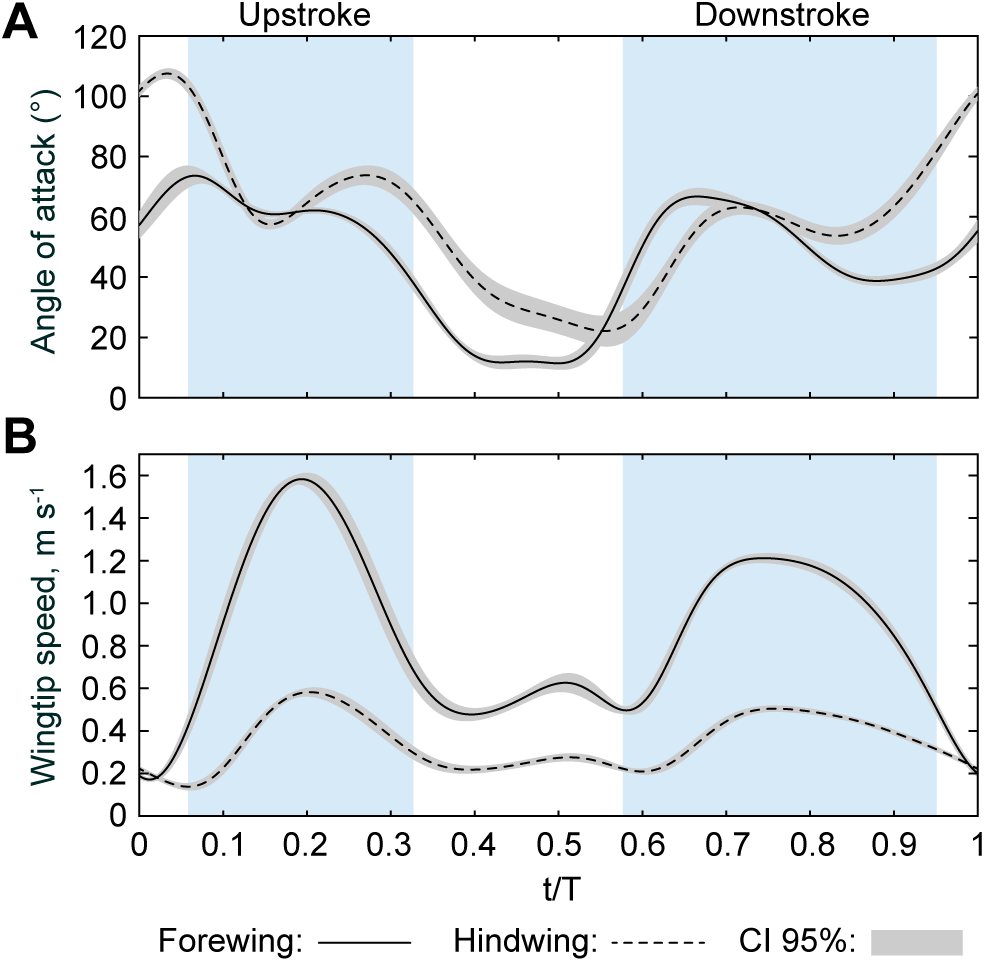
Mean speed at radius of gyration and angle of attack of *Trichogramma telengai* forewings and hindwings аveraged over 11 video sequences.

Stroke plane pitch angle (*β*) is negative in most filmed insects (Fig. 5) and varies in a wide range from −27 to 33°. The body oscillates within the kinematic cycle especially during fling phase (Fig. 6), median body pitch angle (*χ*) amplitude is 12°.

**Figure 5.**
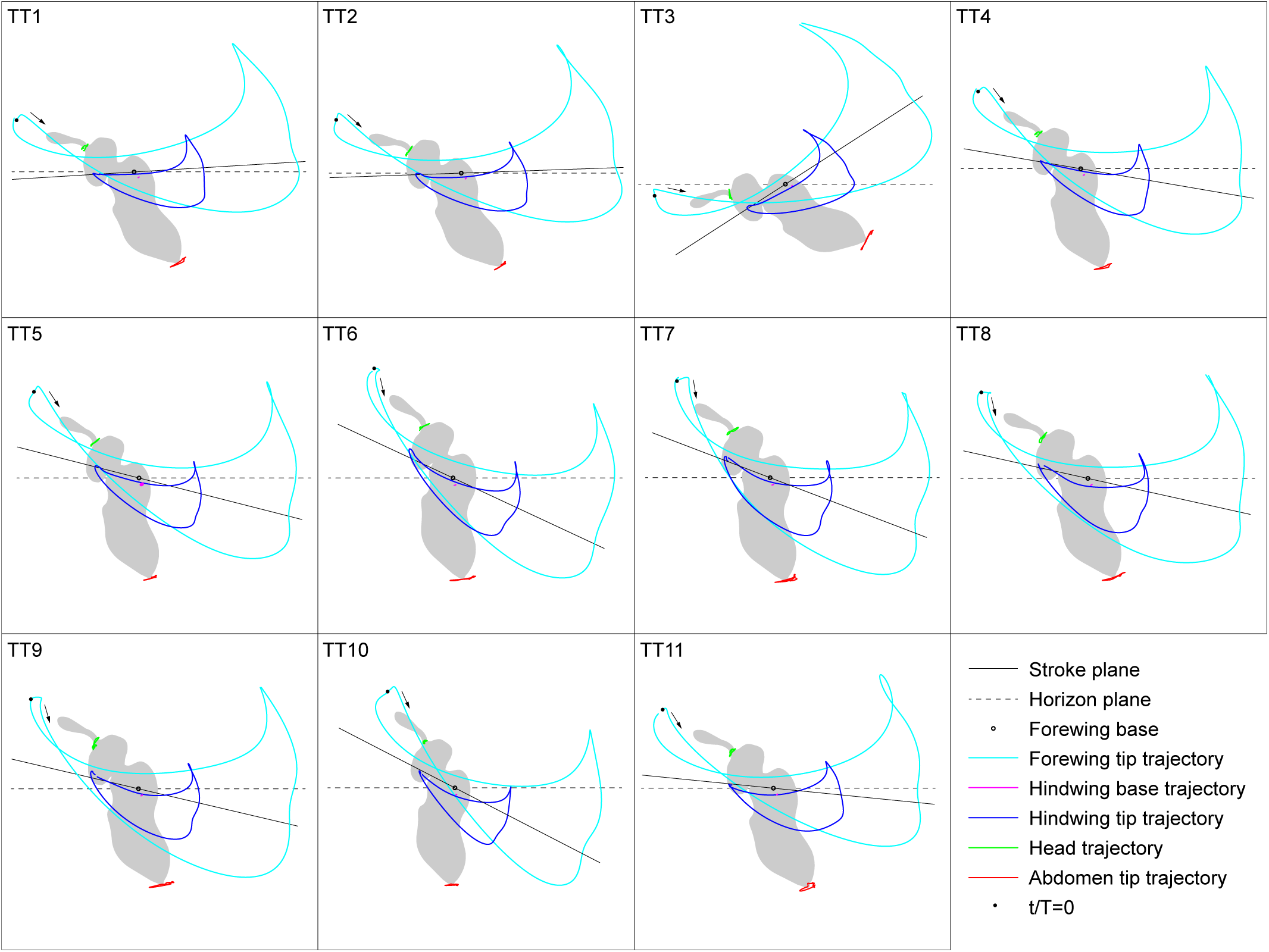
Wing tip trajectories and body orientation of *Trichogramma telengai* specimens TT1-TT11.

**Figure 6.**
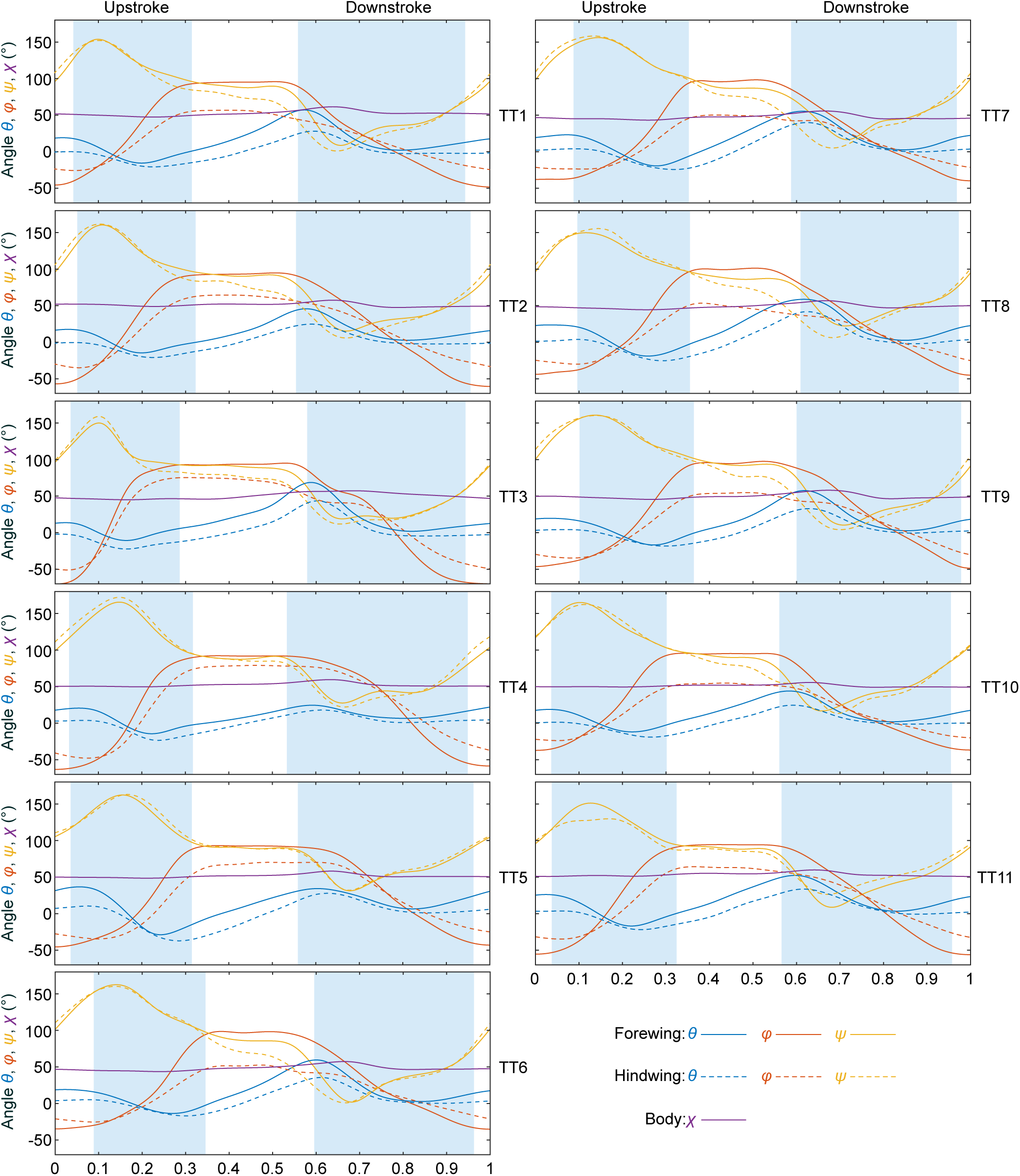
Kinematics description of *Trichogramma telengai* individuals: Euler angles of forewings and hindwings and body pitch angle.

## Discussion

### Body size and wing morphology of *T. telengai*

The mean body mass of *T. telengai* is 4 times less than mass of *E. formosa* (25 µg) (Weis-Fogh, 1973) and three orders of magnitude less than the mass of medium-sized insects like *Musca domestica* (about 6800 µg) (Belyaev and Farisenkov, 2019), two orders of magnitude less than the mass of *Drosophila melanogaster* (about 600 µg) (Burggren et al., 2017) and multiple times greater than the mass of smallest insects like featherwing beetles *Paratuposa placentis* (2.4 µg) (Farisenkov et al., 2022). It is known that *E. formosa* body length is between 0.64 and 0.73 mm (Singh and Sood, 2018), hence *T. telengai* body length is 1.4 times less on average.

Due to the relatively small area occupied by bristles, *T. telengai* wings can be classified as mainly membranous. Bristled area in *E. formosa* is slightly greater (22.7% of forewing and 29.3% of total area) according to our morphometrics based on wings photos provided by Singh and Sood, 2018). Hymenopterans of the Mymaridae family with body size under 500 µm have bristled wings with setae area part greater than 50% (Jones et al., 2016). Similarly, wings of sub-millimeter thrips (Ford et al., 2019) and Ptiliidae beetles (Polilov et al., 2019) are mainly bristled with setae area at least 80 and 90% respectively. Smallest documented flying dipterans such as biting midges *Dasyhelea flaviventris* and gall midges *Anbremia sp.* are larger than a millimeter body size (Lyu et al., 2019). Thus, *T. telengai* represents a model object of smallest membranous-winged insect for flight mechanics investigation.

### Flight performance

Mean and maximal flight speed of *T. telengai* corresponds closely to the values obtained in *Eretmocerus mundus*, miniature aphilinid parasitoid wasp with 0.7 mm body length. Its mean aerial horizontal speed in the absence of wind is 21 and maximal is 25 сm s^-1^ (Sarig and Ribak, 2021). Also, *T. telengai* is close to featherwing beetles in flight performance characteristics (Farisenkov et al., 2020), in particular to *Paratuposa placentis* (body length 0.38 mm). However *Trichogramma* spp. relative flight muscle volume is much greater (Polilov, 2016b) than in ptiliids (Farisenkov et al., 2022), probably because membranous-winged miniature wasps need more body-mass-specific muscle power to achieve flight performance equal to featherwing beetles.

### Features of *T. telengai* kinematics

*T. telengai* (*Re* = 12) demonstrates the same wingbeat cycle type as the smallest previously researched wasp *E. formosa* (*Re* = 17) (Cheng and Sun, 2021): relatively fast U-shaped power strokes at high angles of attack and a slow prolonged clap-and-fling. We can see only minor differences in the kinematics of *E. formosa* and *T. telengai* that could presumably be due to the Reynolds number difference (Fig. 7). *E. formosa* demonstrates slightly but significantly (hereinafter MWU test *p* < 0.05) lower wingbeat amplitude *Δφ* (139°) than *T. telengai*. Recovery stroke is more prolonged in *T. telengai*, and stroke deviation angle (*θ*) reaches larger values at fling phase. Wherein *Δθ* is almost the same in both species (2° difference).

**Figure 7.**
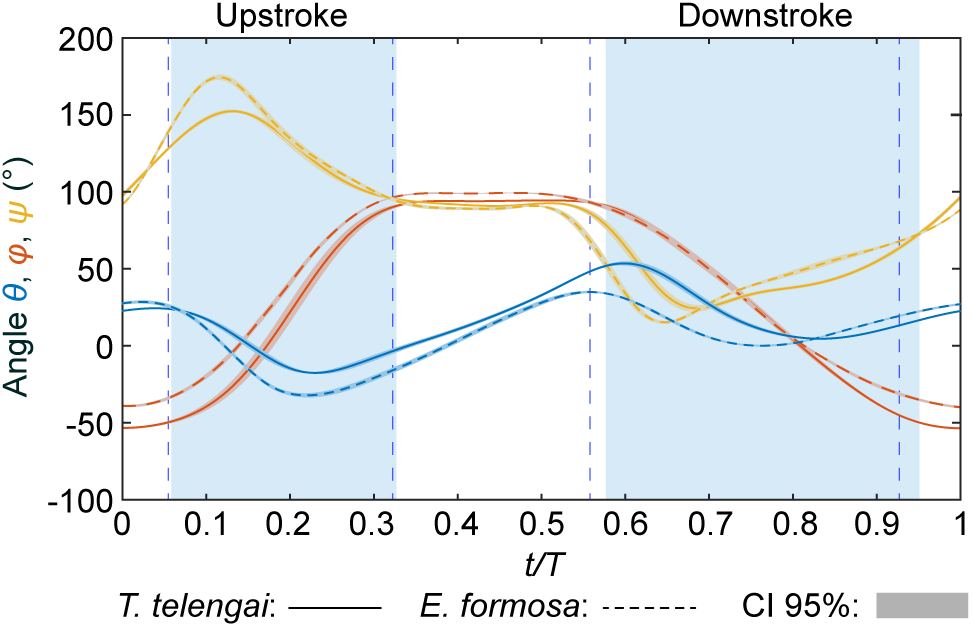
Comparison of *Trichogramma telengai* and *Encarsia formosa* forewing kinematics (Cheng and Sun, 2021): averaged Euler angle time course.

Thrips *Frankliniella occidentalis* (*Re* = 15) (Lyu et al., 2019) have bristled wings (76% of area occupied by setae (Ford et al., 2019)), unlike *E. formosa* and *T. telengai*, but demonstrates nearly the same flight style (Lyu et al., 2019). At *Re* near 10 bristled wings act in the same way as membranous ones, but lessen the muscle power requirements. Two-winged Ptiliidae beetles which are comparable in size (BL = 0.4 mm) to *T. telengai* have significantly modified kinematic cycle: even longer recovery strokes which replace not only the pronation phase, but also the supination. That trajectory of the wings causes high aerodynamic torques and leads to excessive body pitch oscillations (29°) which are compensated by elytra movements (Farisenkov et al., 2022). Body pitch amplitude in *T. telengai* is significant (12°) due to the presence of recovery stroke but not as expressed as in featherwing beetles, so U-shaped wingbeat cycle of microinseсts has less effect on the body in-flight stability.

### Hindwing kinematics

Hindwings are linked with forewings at one point by two hook-like hamuli which are located in the basal half of each hindwing (Fig. 1). Therefore, a hindwing has a significant degree of freedom relative to the forewing and its kinematic parameters differ due to mismatching of wingblade planes and transverse deformations. Hindwing amplitude (***Δ****φ*) is significantly lower: 102° on average (Fig. 6). Hindwing reaches extremes in stroke deviation angle (*θ*) later than forewing (Fig. 6); and alteration in angle of attack demonstrates phase shift in movement of pairs of wings during translations and clap-and-fling (Fig. 4A). A smaller flapping amplitude, a less pronounced plateau of the amplitude angle, and low values of pitch angle indicate that, most likely, the hindwings of *T. telengai* are more deformed during power strokes than the forewings.

## Conclusions

We have shown that the wing kinematics of the miniature parasitoid wasp T. telengai is fundamentally different from the kinematics of large membranous-winged insects and the smallest bristled-winged Ptiliidae beetles. At the same time it is similar to that of the membranous-winged parasitoid wasp E. formosa and bristled-winged thrips F. occidentalis of the same size class. Thus, we assume that there is a small range of Reynolds numbers in which both wing types are sufficiently efficient to perform flight successfully, but flight on membranous wings is more energetically expensive and requires a relatively larger wing muscles volume.

## List of symbols and abbreviations

BL: body length
CI: confidence interval
FPS: frames per second
*J*: advance ratio (body velocity divided by mean wingtip speed)
LED: light-emitting diode
*Re*: cycle-averaged Reynolds number based on the radius of gyration
*R_F_*: forewing length
*R_g_*: radius of gyration
*RgF*: forewing radius of gyration
*R_gH_*: hindwing radius of gyration
*R_H_*: hindwing length
*t/T*: nondimensional time
TT1-TT11: *Trichogramma telengai* specimens
*V*: wing velocity at the radius of gyration
*Α*: angle of attack
*β*: stroke plane pitch angle relative to horizon
*Δφ*: stroke amplitude
*θ*: stroke deviation angle
*ν*: kinematic viscosity of air at 25°С
*φ*: stroke amplitude angle
*χ*: body pitch angle relative to the stroke plane

## Acknowledgements

We are grateful to S.Y. Reznik (Zoological Institute, Russian Academy of Sciences, St. Petersburg, Russia) for providing insects for this study.

## Author contributions

A.A.P. conceptualized and designed this study; S.E.F. designed the experiment and the equipment; S.E.F., N.A.L., E.O.S and A.A.P. collected the data; S.E.F., N.A.L. and D.K. analyzed the data; S.E.F., N.A.L., D.K., E.O.S and A.A.P. wrote the manuscript.

## Data availability

Raw data, extended datasets and Matlab code are avaliable in an OSF repository (https://osf.io/y6t57/?view_only=6293f4addb554f54b8c1a08a28e1cc69)

